# Internal states emerge early during learning of a perceptual decision-making task

**DOI:** 10.1101/2024.11.30.626182

**Authors:** Lenca I. Cuturela, The International Brain Laboratory, Jonathan W. Pillow

## Abstract

Recent work has shown that during perceptual decision-making tasks, animals frequently alternate between different internal states or strategies. However, the question of how or when these emerge during learning remains an important open problem. Does an animal alternate between multiple strategies from the very start of training, or only after extensive exposure to a task? Here we address this question by developing a dynamic latent state model, which we applied to training data from mice learning to perform a visual decision-making task. Remarkably, we found that mice exhibited distinct “engaged” and “biased” states even during early training, with multiple states apparent from the second training session onward. Moreover, our model revealed that the gradual improvement in task performance over the course of training arose from a combination of two factors: (1) increased sensitivity to stimuli across all states; and (2) increased proportion of time spent in a higher-accuracy “engaged” state relative to biased states. These findings highlight the power of our approach for characterizing the temporal evolution of multiple strategies across learning.

## Introduction

Perceptual decision-making is an essential component of human and animal behavior that requires integrating sensory information and selecting an appropriate action (O’Connell et al., 2018). For example, we see a traffic light turn red and make the decision to apply the brake. Scientists have studied perceptual decision-making using carefully designed tasks in which animals learn to identify task-relevant sensory cues while ignoring the task-irrelevant variables (Gold & Shadlen, 2007; Hanks & Summerfield, 2017; International Brain Laboratory et al., 2021). Classic models of sensory decision-making, such as signal detection theory (Green & Swets, 1966; Klein, 2001) or the drift-diffusion model (Ratcliff & Rouder, 1998; Ratcliff & McKoon, 2008), assume that animals use a consistent decision-making strategy across time. However, recent work has shown that animals switch between multiple distinct strategies even within a single session (Ashwood et al., 2022; Bolkan et al., 2022; Hulsey et al., 2024; Weilnhammer et al., 2021).

This previous work identified multiple strategies in mice performing perceptual decision-making tasks by using a Hidden Markov Model (HMM) with Generalized Linear Model (GLM) observations (Ashwood et al., 2022; Bolkan et al., 2022). The resulting “GLM-HMM” provides a powerful framework for identifying internal states during sensory-driven behavior that are not directly observable but are known to modulate behavior nevertheless (Flavell et al., 2022; Calhoun et al., 2019). In this model, each hidden state corresponds to a distinct decision-making strategy, parametrized by a set of GLM weights that describe how the animal weighs different task covariates in order to make a decision in that state. When applied to visual decision-making data collected by the International Brain Laboratory (IBL), this method revealed that fully trained mice switch between three distinct states that tend to persist for many trials in a row: an “engaged” state in which performance is high, and two “biased” states, in which performance is lower, with left and right biases, respectively (International Brain Laboratory et al., 2021; Ashwood et al., 2022). However, it remains unknown whether these distinct states or strategies are present early in training, or how they evolve over the course of learning. The GLM-HMM is unfortunately ill-equipped to address this question, given that it captures abrupt changes between discrete states but does not allow for gradual changes in parameters or states over time.

To overcome this limitation, we introduce “dynamic GLM-HMM”, an extension of the GLM-HMM in which the model parameters are allowed to evolve across sessions. Our model can be used to characterize state-dependent decision-making behavior that varies across sessions. Specifically, this model allows us to examine whether mice rely on a single or multiple strategies during training, how exactly these strategies evolve over learning, and whether animals change how much time they spend in each state. Our approach draws inspiration from the Psytrack model (Roy et al., 2021), which consists of a dynamic GLM with time-varying weights but does not contain multiple states. Note that our method provides a simpler, more interpretable approach to recent work from (Bruijns et al., 2023), which uses an infinite semi-Markov model to examine time-varying multi-state choice behavior.

We apply the dynamic GLM-HMM to a training dataset of mice learning to perform a visual decision-making task (International Brain Laboratory et al., 2021). Remarkably, we show that mice switch between distinct “engaged” and “biased” states since the beginning of training. For most mice, a three-state model outperforms a single-state model in accounting for decision-making behavior as early as the second session. Moreover, we show that mice improve their performance over the course of training not only because of increased sensitivity to stimuli, predominantly in the engaged state, but also because they spend an increasingly larger fraction of time in the engaged, higher-accuracy state. Finally, we use the dynamic GLM-HMM to propose a novel criterion for determining when an animal has successfully learned the task; namely, when it achieves a certain level of accuracy in the engaged state. We show that by this criterion, mice have often learned the task earlier than indicated by conventional measures like percent correct, which do not take internal states like disengagement into account.

## Results

### Characterizing multiple time-varying strategies with dynamic GLM-HMM

The standard GLM-HMM consists of a hidden Markov model (HMM) with multiple latent states, where each state corresponds to a distinct Bernoulli generalized linear model (GLM) governing the mapping from sensory inputs to the choices (Figure 1A,C). The dynamic GLM-HMM extends this model by allowing both the HMM transition matrix, which governs the probability of switching between states, and the GLM weights, which define the different decision-making strategies, to vary across sessions. Specifically, we allow the GLM weights to change from session to session according to a Gaussian prior (Figure 1B), and we allow per-session HMM transition probabilities to vary according to a global Dirichlet prior (see Methods). Taken together, these extensions allow decision-making strategies to change gradually across sessions, and for the transitions between them to vary so that different strategies are employed more or less frequently in different sessions.

**Figure 1:**
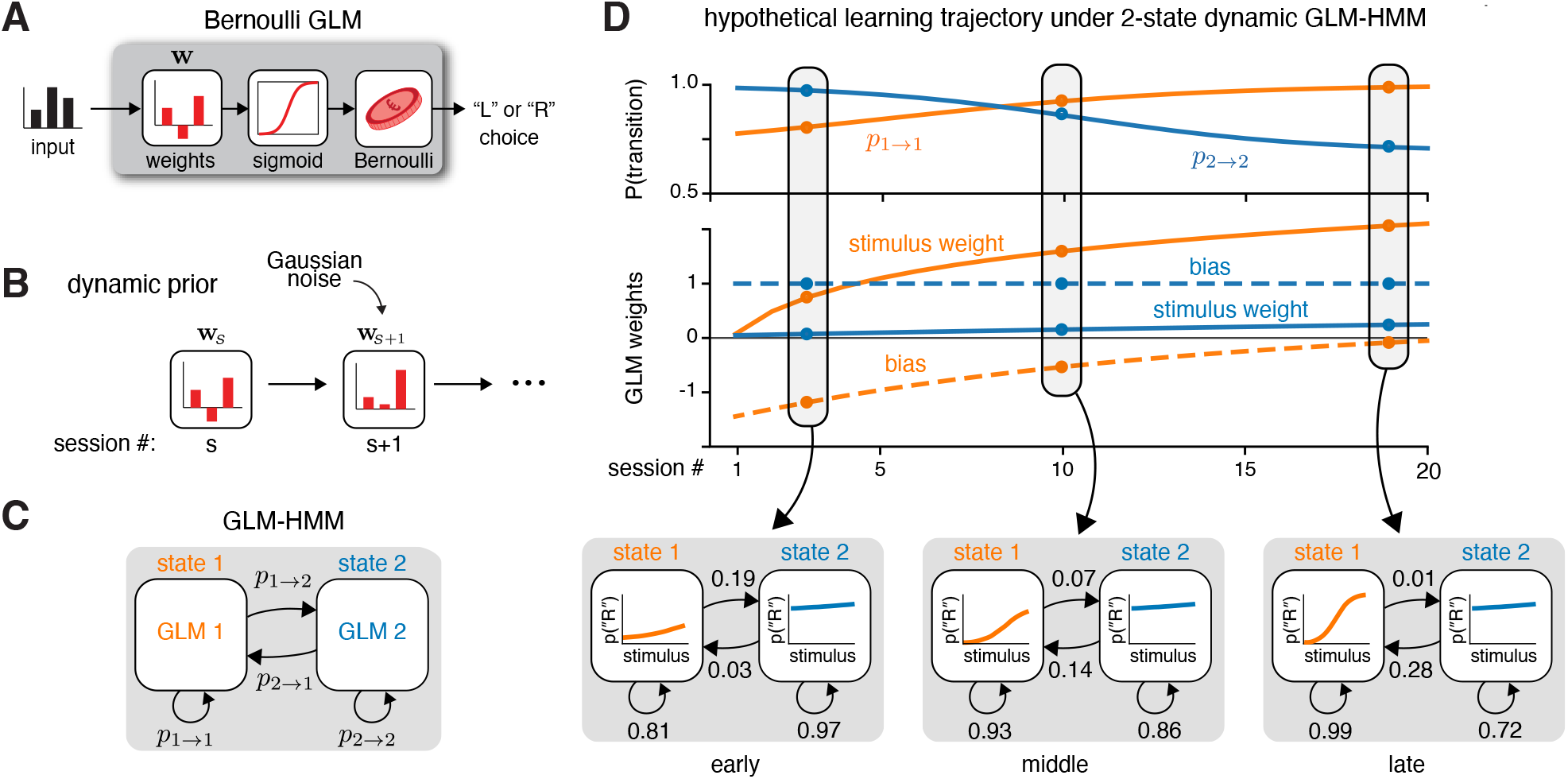
Characterizing multi-state learning trajectories with dynamic GLM-HMM. **(A)** Schematic of the single-state Bernoulli generalized linear model (GLM). The model takes in a vector inputs on each trial (e.g., stimulus contrast, previous choice). This input vector is linearly combined with a weight vector **w**, then transformed by a logistic function to produce the probability for a Bernoulli random variable, which determines the binary choice (left or right) on each trial. **(B)** To allow changes in GLM weights across sessions, we use a Gaussian prior centered on the weights from the previous session. The variance of this Gaussian controls how much the GLM weights can change between sessions. **(C)** Schematic of a 2-state GLM-HMM, a Hidden Markov Model parametrized by a set of probabilities governing the transitions between states, and a pair of Bernoulli GLMs governing outputs in each state. **(D)** Hypothetical trajectories of a set of self-transition probabilities (above) and GLM weights (middle) for a two-state dynamic GLM-HMM during learning. The self-transition probabilities *p*_1_ →_1_ and *p*_2_ →_2_ describe the probability of remaining in state 1 or state 2 after a trial, also known as ‘stickiness’ of each state. The GLMs contain two weights governing the animal’s decision-making strategy in each state: a sensory stimulus weight and a bias weight. Here, the state 1 weights evolve to have increased stimulus sensitivity and decreased bias over time, while the state 2 weights mostly remain constant. Dots represent the model parameters for 3 example sessions from “early”, “middle”, and “late” training sessions, which correspond to the psychometric curves and transition probabilities shown below.

The dynamic GLM-HMM contains hyperparameters that govern the amount of change in parameters across sessions, allowing the model to capture both slow and rapid rates of learning. The fitted weights can range from completely static, in which animals use a fixed set of strategies across all sessions, to highly variable, in which animals use a nearly independent set of strategies on each session. Similarly, the fitted transition matrices can range from completely static, in which animals switch between states at fixed probabilities across the sessions, to highly variable, in which animals rapidly change their switching behavior at different sessions during training. We validated our novel method by first fitting on simulated datasets, succesfully recovering true parameter trajectories and true hyperparameters used to generate the data (Supplemental Figure 1).

Within our framework, task learning corresponds to an evolution of the GLM weights and transition probabilities corresponding to an increase in performance. Figure 1D shows a hypothetical learning trajectory in a perceptual decision-making task. In this schematic, the imagined animal is assumed to have two strategies, each governed by a different GLM with two weights: a bias weight and a weight on the sensory stimulus. We can use these weights to compute instantaneous psychometric curves, which describe the probability of a rightward choice as a function of the stimulus strength for each session individually.

In early sessions, the sensory weights in both states are close to 0, corresponding to flat psychometric curves and low performance (Figure 1D). Moreover, states 1 and 2 initially exhibit left and right biases, respectively, given by non-zero bias weights of opposite signs. Over the course of training, the sensory weight in state 1 (in orange) gradually increases, while the bias weight decays toward 0. This suggests that the hypothetical animal is learning to solve the task as its choices in this state rely increasingly more on the sensory information and increasingly less on a left-side bias. Accordingly, the psychometric curve associated to state 1 attains in late sessions the characteristic “S-shape” that corresponds to high performance (Figure 1D). Thus, state 1 represents a “task-engaged” or “engaged” state. In contrast, state 2 represents a “right-biased” state in which the animal performs sub-optimally since its bias weight is consistently high throughout all sessions and its stimulus weight is close to 0.

In this schematic, the self-transition probabilities are generally high, suggesting that the states tend to persist for many consecutive trials (Figure 1D). In early sessions, state 2 has higher self-transition probabilities than state 1, whereas in late sessions the opposite holds. This means that state 1, the engaged state, becomes more persistent as sessions progress. As a result, the time spent being in the engaged state also increases over sessions, leading to an improvement in the overall task performance. In this example, the imagined animal employs both strategies since the very first training session.

A key question our work addresses is whether multiple strategies arise only after an animal receives extensive exposure to the task. To rigorously answer this question, we first ensure with another simulated dataset that our model can also characterize a situation in which the number of strategies used is flexibly changing over time. We simulate a dataset in which an imagined animal only uses a single state during the first 10 sessions (state 1) and begins alternating between all 3 states later on (Supplemental Figure 2). Although dynamic GLM-HMM uses a fixed number of total states, we showed that our method can characterize well behavior that employs different numbers of states at different sessions. Crucially, we used a similar number of simulated trials as in a real perceptual decision-making experiment, thus proving that given a realistic amount of data our method is flexible enough to uncover states that are not necessarily used across all sessions.

### Choice behavior of example mouse is well explained by three dynamic strategies

To examine how animals employ multiple time-varying strategies during learning, we fit the dynamic GLM-HMM to visual decision-making data collected from 32 mice in 3 different labs within the International Brain Laboratory (International Brain Laboratory et al., 2021). In this 2-alternative forced-choice task, mice were presented with a sinusoidal grating on the left or right side of the screen, and had to report its side by turning a wheel. Task difficulty was controlled by varying the contrast of the grating between 0% and 100% (Figure 2A) (International Brain Laboratory et al., 2021). Note that mice were trained on the task in two stages: first on a basic task with equal left-right stimulus probability (50:50), and then on a full task where stimulus probabilities alternated between 20:80 and 80:20 across trial blocks with variable durations (International Brain Laboratory et al., 2021; Burgess et al., 2017).

**Figure 2:**
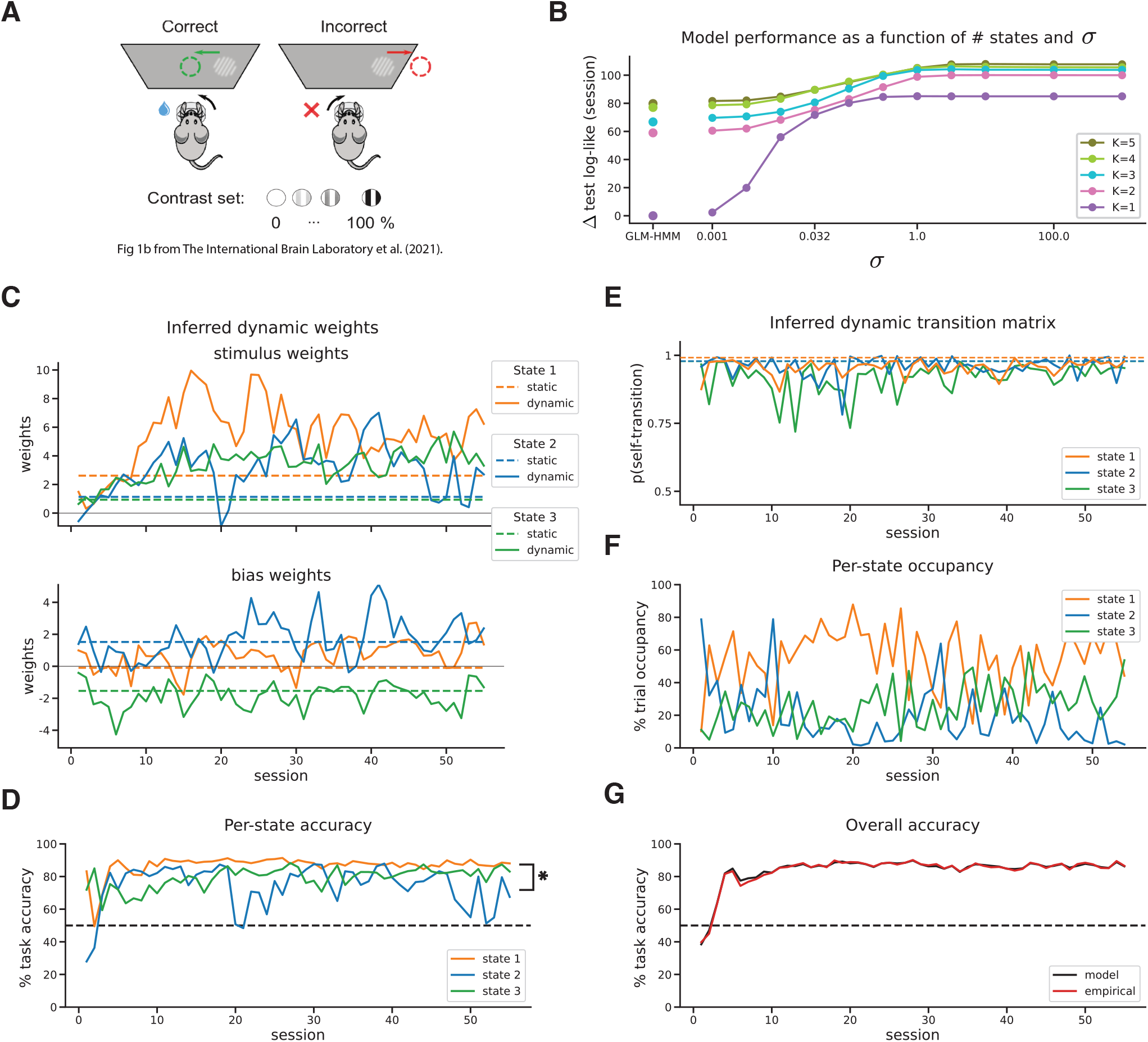
Choice behavior of example mouse is well explained by three dynamic strategies. **A**. Schematic for visual decision-making task (adapted from International Brain Laboratory et al. (2021)). **B**. Model performance comparison for example animal (ibl witten 012) between dynamic GLM-HMMs with different numbers of latent states *K* and rates of change of the weights *σ*. Model performance is measured as the average delta test log-likelihood per session. **C**. Inferred dynamic GLM weights for the best fitting 3-state model (*σ ≈* 3) for two task variables, signed stimulus contrast (top) and bias (bottom), during the first 45 sessions. Inferred weights for the standard GLM-HMM are shown with dashed lines. **D**. Per-state task accuracy % across sessions, computed using the inferred weights from C. **E**. Inferred dynamic transition matrix for the best fitting 3-state model (*α =* 2). Inferred probabilities for the standard GLM-HMM are shown with dashed lines. **F**. Per-state trial occupancy % across sessions, computed by hard assigning the latents based on the highest posterior probability. **G**. Overall model and empirical task accuracy % across sessions.

We fit the model independently to the behavioral data from each mouse, using all trials from the training (basic task) and post-training periods (full task). We modeled animals’ choices using four task inputs: the signed stimulus contrast (positive for the right side and negative for the left side), an offset or bias, the animal’s choice on the previous trial, and the rewarded side of the previous trial. To closer match the animal’s perception of the visual stimulus, we used a tanh transformation of the stimulus contrast that better explained the choice data than no transformation at all (Supplemental Figure 3E), in line with previous work (Roy et al., 2021). Note that for this task the optimal strategy that maximizes reward corresponds to a large positive weight on the signed stimulus contrast and zero weights on all other inputs, which means no dependence on trial history or bias.

Figure 2 and Figure 3 show results for an example mouse, where the first 10 sessions correspond to the basic task that has equal left-right stimulus probability. To determine how many discrete strategies the animal used and how variable in time these strategies are, we compared performance across models with 1, 2, 3, 4, and 5 states, respectively, and with different values of the hyperparameter *σ* governing the evolution of all the weights (Figure 2B). The models performed best for *σ* ≈3, which corresponds to an intermediate rate of change of the weights. We found that three states are sufficient to attain high performance on the test set, in line with previous work that used three states to characterize the IBL decision-making data in fully trained animals (Ashwood et al., 2022).

**Figure 3:**
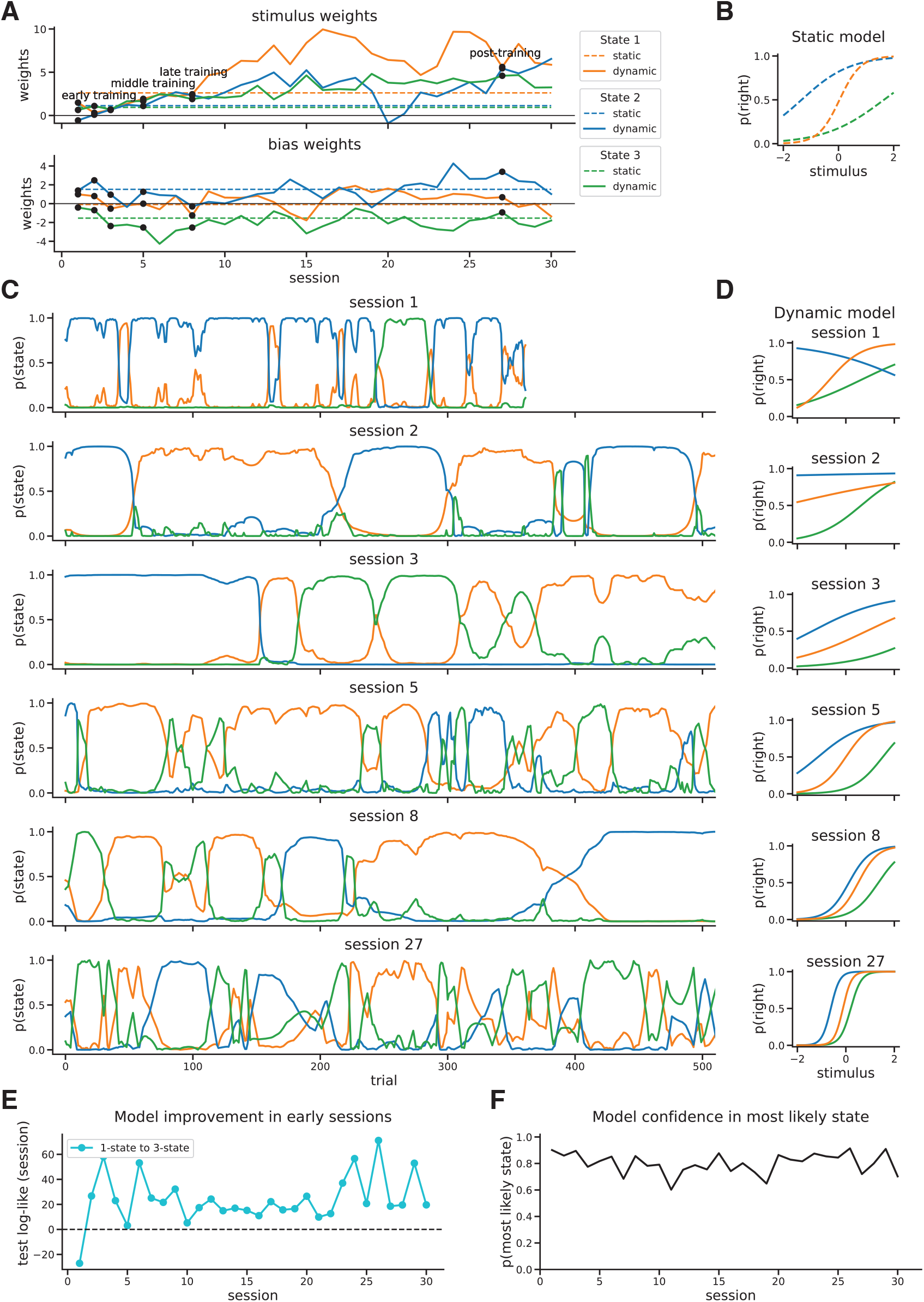
Example mouse employs multiple strategies since the beginning of training. **A**. Stimulus and bias weights for the best three-state models for example mouse. Black dots are the chosen example sessions. **B**. Perstate psychometric curves for the static model. **C**. Posterior probabilities of the latent states on a trial-by-trial basis for example sessions from different training epochs. **D**. Per-state psychometric curves for the example sessions from C. **E**. Difference in average test log-likelihood per session between best 3-state and 1-state dynamic models. **F**. Average posterior probability of most likely state for each session.

Note that the standard GLM-HMM was consistently outperformed by a dynamic GLM-HMM, which shows that static weights are not appropriate for fitting training data (Figure 2B). Interestingly, there was a substantial increase in performance from 3 states to 4 states in the standard GLM-HMM that was not present in the dynamic GLM-HMM. Since standard GLM-HMM uses static weights, the model will often overestimate the number of states needed for capturing time-varying strategies – a limitation that dynamic GLM-HMM surpasses.

The decision-making strategies associated to the latent states were mainly distinguished by the stimulus and bias weights, since the other weights were close to 0 across all states and sessions (Supplemental Figure 4A). The stimulus weight in state 1 (in orange) increased rapidly over the first 10-20 sessions, reflecting the learning trajectory of this animal as it was relying on the stimulus information increasingly more as the training progressed (Figure 2C). The bias weights in state 1 remain around 0 throughout, suggesting that the animal generally did not have a biased preference in this state. Thus, state 1 represents an “engaged state”, since the task accuracy and stimulus weight in this state increased during training and remained consistently high after that (Figure 2C,D). Moreover, the animal’s performance in state 1 was significantly higher than its performance in the other two states (Figure 2D).

States 2 and 3 are characterized by large bias weights with opposite signs, which correspond to right and left biases, respectively. Surprisingly, states 2 and 3 also showed a substantial increase in the stimulus weights over sessions, suggesting that the animal was also learning in these other states. This finding was also supported by the increase in task accuracy over time in all three states, all surpassing chance performance at 50% during most sessions (Figure 2D). Thus, state 2 and state 3 represent “right-biased” and “left-biased” states, respectively, with partial task engagement that fluctuated across sessions. Note that the overall model accuracy computed using the three per-state accuracies closely matched the animal’s empirical task performance, an additional validation of our modeling procedure (Figure 2G).

The inferred self-transition probabilities were generally high, showing some variability and lower values at earlier sessions (Figure 2E). In all, the states are persistent for many consecutive trials during the majority of sessions, and we did not observe any systematic change in the transition probabilities over time.

We next examined the posterior probabilities of the latent states, i.e., the per-trial likelihood of the animal being in a particular state after the model is fit to data. Visualizing example sessions revealed how this animal switched between states across different training epochs during early, middle, late, and post-training (Figure 3A,C). Although sometimes the animal switched quickly between states, the animal generally spent tens to hundreds of consecutive trials in the same state (Figure 3C). For these example sessions, the different behavioral strategies are seen in the psychometric curves, which represent the probabilities of a rightward choice as a function of the signed contrast of the visual stimulus (Figure 3D).

In the first few sessions, the psychometric curves were quite flat since the stimulus weights and task performance were low at the beginning of training across all states (Figure 3D, Figure 2C,D). Over the course of training, the psychometric curves, especially for the engaged state, became increasingly steeper, in line with the increase in the stimulus weights over sessions that reflect the animal’s learning of the task (Figure 3A,D). The biases towards rightward and leftward choices in states 2 and 3, respectively, are apparent in the psychometric curves of these two states (Figure 3D). As expected, the psychometric curves in the post-training example session were similar to the psychometrics from the static model (Figure 3B,D), with the engaged state attaining the characteristic “S-shape” curve centered at 0 that corresponds to high task performance.

The trial occupancies for all states were consistently higher than 0, meaning that this mouse used all strategies in the majority of sessions (Figure 2F). The probabilities of the latent states in Figure 3C show that even in the first few sessions, the mouse spent time in all three states. Moreover, we can be very confident about which state the animal was in since the likelihood of the most probable state was around 0.7-0.9 throughout most sessions (Figure 3F). Remarkably, the 3-state model outperformed the single-state one for all sessions starting with the second one (Figure 3E). Taken together, these findings suggest that there were multiple strategies present in this example mouse’s behavior since the very beginning of training.

### State-specific strategies are similar across mice and present since the start of training

Next, we examined the dynamic GLM-HMM fits across all 32 mice to understand how the previous findings with regards to the example animal generalize across all animals. When fitting our model to data from each animal individually, we initialized the model with the fitted parameters from the global standard GLM-HMM, which used the data from all mice together (see Methods). We found that the results above for the example mouse are consistent across all mice.

Plotting average model performance across animals as a function of the weights’ rate of change *σ* reveals that models with *σ* ≈ 3 performed much better than models with static weights (Figure 4A). The 3-state dynamic GLM-HMMs with *σ ≈* 3 attained close to maximum log-likelihood on the test set, suggesting that three states are sufficient to describe most animals’ choices (Figure 4A,B), in line with previous work (Ashwood et al., 2022). Therefore, we chose the 3-state model for each animal for further analysis.

**Figure 4:**
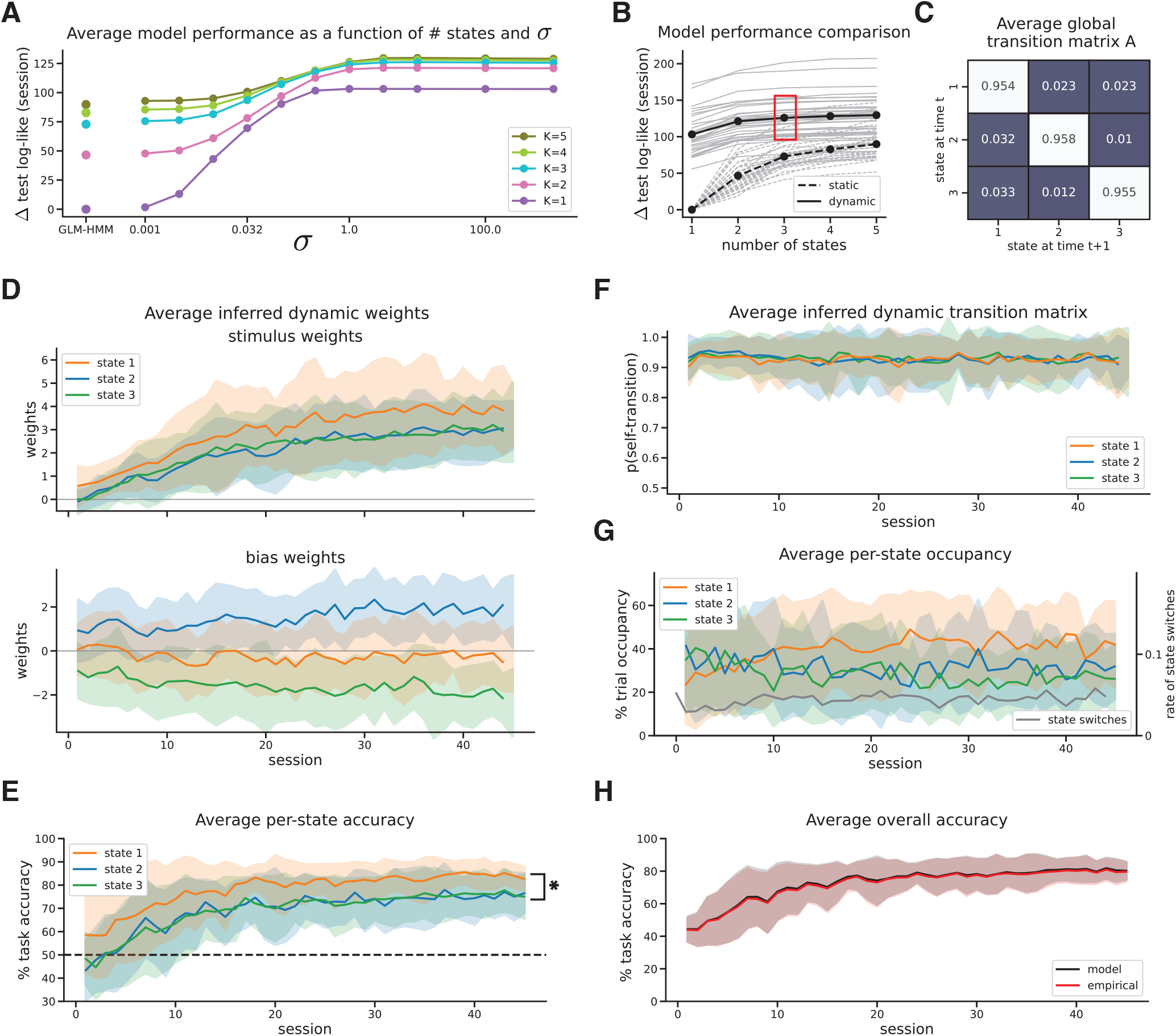
Choice behavior of all mice is well explained by three dynamic strategies. **A**. Δ test log-likelihood (averaged across sessions) as a function of hyperparameter *σ* for models with 1, 2, 3, 4, and 5 states, respectively, averaged across all animals. **B**. Δ test log-likelihood as a function of number of states, averaged across animals (in black) for models with optimal hyperparamter *σ =* 3.2, or static models. Individual lines are shown in gray for each animal. **C**. Average global transition matrix across animals for best fitting 3-state models **D**. Average inferred dynamic GLM weights for the best 3-state models (*σ =* 3.2) for two task variables, stimulus (top) and bias (bottom), shaded area is −/ + 1 STD across animals. **E**. Percentage task accuracy for each state and session, averaged across animals (shaded area is −/ + 1 STD). State-1 accuracy is significantly higher than mean accuracy in states 2 and 3 for each animal across sessions **F**. Average inferred dynamic self-transition probabilities for the best 3-state models for each session (*α* = 2), shaded area is −/ + 1 STD across animals. **G**. Percentage trial occupancy for each state and session, averaged across animals (−/ + 1 STD in shaded area) (left axis). Rate of state switches as a function of session number (right axis). **H**. Average model and empirical % task accuracy for each session, averaged across animals (1 STD in shaded area).

The behavioral strategies associated to the latent states were distinguished by the stimulus and bias weights, since the history-dependent weights were on average across animals close to 0 (Supplemental Figure 4B). The inferred weights showed some variability across animals (Figure 4D), but nevertheless embodied consistent strategies across all mice. The stimulus weights in state 1 (in orange) were largest, whereas the rest of the weights in this state were close to 0. Thus, state one represents an engaged, unbiased state, in which the animals eventually acquired high task performance (Figure 4D, E).

Similar to the example mouse, states 2 and 3 represent partially engaged right-biased and left-biased states, respectively, in which animals had significantly lower task performance than in the engaged state 1 (Figure 4D, E, see Methods, Significance Testing). Note that the bias weights in state 2 and state 3 increased in absolute value over time, suggesting that the animals were becoming increasingly more biased in their choices when they were using those two strategies (Figure 4D). Importantly, the overall model task accuracy computed using the per-state accuracies closely matched the empirical task performance across animals (Figure 4H).

In line with previous work (Ashwood et al., 2022), the average global transition matrix across animals for the best-fitting 3-state models has large elements on the diagonal, meaning that the animals were very likely to persist in the same state for many trials in a row (Figure 4C). For the dynamic transition matrix, the self-transition probabilities of all states were on average close to 0.8, confirming the persistence of the states on a session by session basis. Although there was no systematic change over sessions in the self-transition probabilities when averaged across animals, there was a lot of variation for individual animals across sessions (Figure 4F). This finding was supported by obtaining a low optimal hyperparameter *α* for the per-session transition matrix prior, which means that the best fitting models allowed transition probabilities to change abruptly from one session to another, even though that happened rarely (Supplemental Figure 3D). The rate of switching between states remained the same across all sessions (Figure 4G).

On average across animals, the time spent in each state was higher than 20% for all states and all sessions, suggesting that the animals used all three behavioral strategies during all stages of training (Figure 4G). Moreover, the model was highly confident when inferring the state the animals were in since the most likely latent state had the average probability at around 0.8 for all sessions, including early ones (Figure 5B).

**Figure 5:**
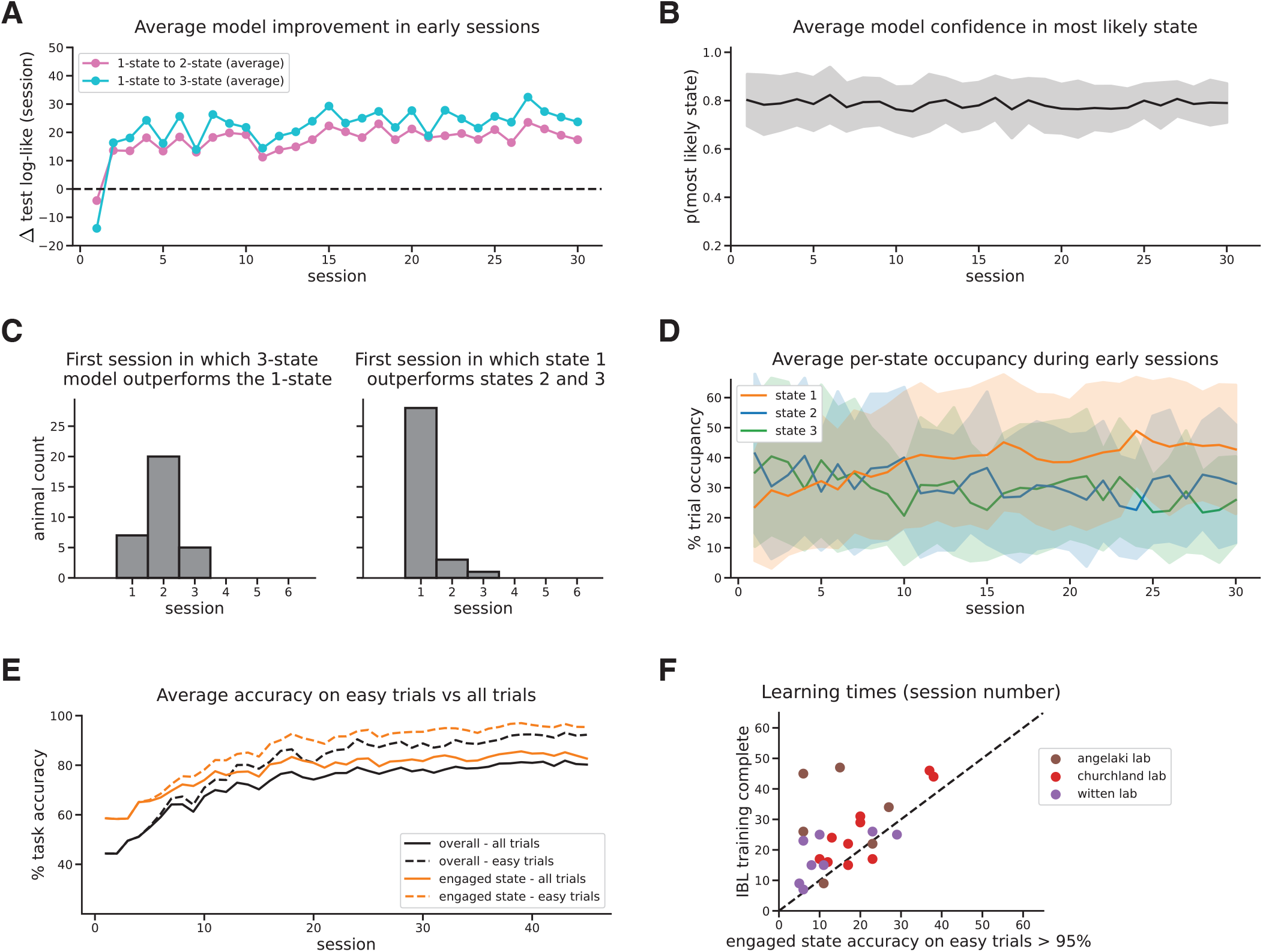
Internal states in mice emerge early during learning. **A**. Difference in average test log-likelihood per session between best 3-state and 1-state models (in blue) and best 2-state and 1-state models (in pink), averaged across animals and shown for the first 30 sessions. **B**. Average posterior probability of most likely state for each session, averaged across animals (shaded area is −/ + 1 STD). **C**. Histogram across animals for first session in which 3-state model performs better than 1-state one (left). Histogram across animals for first session in which state 1 (engaged state) performs better than the other two states (right). **D**. Percentage trial occupancy for each state during the first 30 sessions, averaged across animals (−/ + 1 STD). **E**. Percentage task accuracy in the engaged state and overall during easy trials versus all trials, averaged across animals. **F**. Scatter plot of learning times based on our criterion (engaged-state accuracy reaches 95% on easy trials) versus the IBL definition of the end of training, across animals from the three labs chosen (Laboratory, 2022).

To further confirm that three distinct strategies best explain the animals’ choices even during early training, we plotted for each session the average difference in test log-likelihood between the best fitting 3-state and 1-state dynamic models (Figure 5A). Remarkably, we found that the 3-state model outperformed the 1-state one for every session except the first, and for the majority of animals this held starting as early as the second session (Figure 5A, C). In terms of the amount of correct predictions of animals’ choices, the 3-state dynamic model had on average 5% more correct predictions than the 1-state one (Supplemental Figure 3B).

In all, our findings show that these internal states were present and identifiable even in the early training periods, suggesting that the animals’ biased strategies did not arise due to extensive exposure to the task. In particular, the animals had distinct right-biased and left-biased strategies even when the left-right stimulus probability was 50-50%, before the biased blocks were introduced.

Animals learned to solve the task within the first 10-20 sessions, as both their task accuracy and stimulus weights in state 1 increased (Figure 4D,E,H). Surprisingly, states 2 and 3 also showed an increase in the stimulus weights over sessions, suggesting that the animals were also learning in these other states, albeit less than in the engaged state. To verify that indeed learning happens in all states, we built a control model where we only modify the stimulus weights in states 2 and 3 to be static and found that the control model fits the data significantly worse (Supplemental Figure 3C). For the majority of animals, the earliest session in which the engaged state (state 1) had higher accuracy than the other states is the first session, confirming that the identity of the states were consistent even during early training (Figure 5C).

Interestingly, the time animals spent in the engaged state significantly increased over the first 15 sessions (Figure 5D, see Methods, Significance Testing). Early on, the animals were spending more time in the biased states, whereas later on they were spending more time in the engaged state. Therefore, animals improved their accuracy on the task through a combination of two changes: the stimulus weights grew larger, especially in the engaged state, and the animals spent increasingly more time in the engaged state.

On easy trials, defined as trials in which the stimulus contrast was at least 50%, the task accuracy in the engaged state increased rapidly and plateaued close to 100% (Figure 5E). To determine an animal’s end of training within our framework, we defined the time it takes to learn the task as the first session in which performance in the engaged state reaches 95%. We chose to formalize learning based on the engaged-state accuracy rather than the overall one because an animal could perform poorly overall if they spent most of the time in the biased or disengaged states, even though they had learned the contingencies of the task and performed very well in the engaged state.

Using this new criterion, we found that the animals succeeded in learning the task at variable times (Figure 5F), in line with previous findings (International Brain Laboratory et al., 2021). However, our criterion led to consistently earlier learning times across animals when compared to the learning times defined by IBL, which were mainly determined by the first session when an animal reaches 90% overall task accuracy on easy trials (Figure 5F). We posit that the IBL’s criterion might overestimate the time it takes an animal to learn the contingencies of the task since they do not account for the animal’s disengagement. However, IBL’s criterion may be more practical, since we generally need fully trained animals to not only understand the contingencies of the task but also to remain engaged.

Lastly, we investigated the dynamics of the inferred states during the post-training epoch, when biased blocks were introduced. During a biased block, the probability of the stimulus appearing on one “biased” side is 0.8 for many consecutive trials, and the task structure alternates multiple times within the same session between right-biased blocks and left-biased blocks (Figure 6A). Interestingly, we observed that some animals performed better on one type of biased blocks compared to the other (Figure 6B).

**Figure 6:**
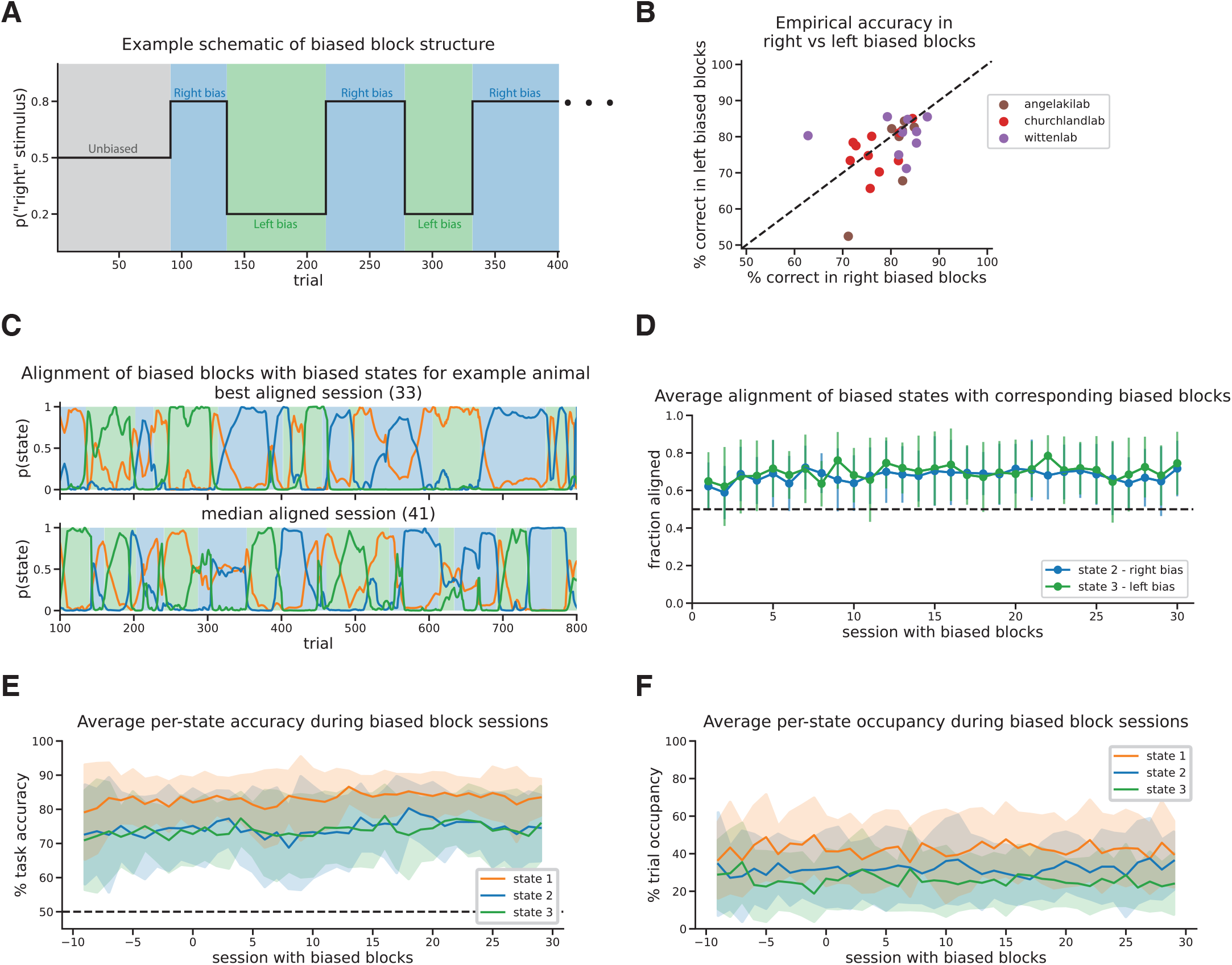
Mice align their biased strategies with the task’s biased blocks. **A**. Schematic of biased block structure in every session during the post-training epoch (blue for right bias, green for left bias). **B**. Scatter plot of % correct responses in right biased blocks vs % correct responses in left biased blocks across animals in 3 labs. **C**. Posterior probability of the latent states for an example animal during two sessions, with biased block structure shown in the background. Sessions represent the best aligned session (top) and median aligned session (bottom) based on alignment computed in D for this example animal. **D**. For states 2 (right-bias) and 3 (left-bias), the average fraction of trials aligned with corresponding biased blocks out of total trials of the time spent in that state across all blocks. Fraction alignment for individual animals are shown in the background. Session 0 represents the first biased block session for each animal. **E**. Per-state task accuracy %, averaged across animals (shaded area is −/ + 1 STD) and aligned to the first biased block session for each animal (session 0) **F**. Per-state task occupancy %, averaged across animals (shaded area is −/ + 1 STD) and aligned to the first biased block session for each animal (session 0).

We also examined the degree to which an animal’s internal state aligned with the experimental structure, and found that animals, when using a biased behavioral strategy, spent on average more time in the corresponding biased block compared to the opposite one (Figure 6D). This is consistent with previous findings showing a substantial degree of alignment between the biased blocks and the animal’s side bias, which was modeled as fluctuating on a trial-by-trial basis (Roy et al., 2021).

Figure 6C shows for an example animal the sessions that had the highest alignment between biased states and same-biased blocks (top) and the median alignment (bottom), respectively. Thus, the animals seemed to adapt to the post-training task structure by reusing a biased strategy that did not require full task engagement but nevertheless led to good performance during the corresponding biased blocks. Note that this alignment seemed to change with experience, since there was a slight increase over time in the fraction of aligned trials and in the biased-states task performance during the post-training epoch (Figure 6D,E). There was no systematic change observed in the time spent in each state after the biased blocks were introduced (Figure 6F).

## Discussion

In this work, we developed the dynamic GLM-HMM, a model for state-dependent perceptual decisionmaking with time-varying weights and state transition probabilities. Unlike the static GLM-HMM, this model is well-suited to nonstationary behavior such as learning, since it allows strategies and their probabilities of occurrence to change over time as an animal learns. We applied this method to a large behavioral dataset of mice learning to perform a sensory decision-making task, in which mice typically started near chance performance and gradually improved their accuracy over the course of many training sessions.

Consistent with previous work (Ashwood et al., 2022), we found that mouse decision-making behavior could be well described by a 3-state model, with a higher-accuracy “engaged” state and two lower-accuracy “biased-right” and “biased-left” states. More significantly, we found that these states could be reliably identified early in training; in all mice, the 3-state model outperfomed the 1-state model within the first three training sessions. The three strategies identified by the model evolved smoothly over the course of training as animals learned to rely more strongly on sensory information when making a decision. Although the rate of learning varied substantially across animals, we found that performance improved as a result of an increase in the sensory stimulus weight in all three states, as well as an increase in the amount of time spent in the engaged, high-accuracy state.

### Relationship to previous work

The dynamic GLM-HMM can be viewed as a straightforward synthesis of Psytrack, a model for continuously varying psychophysical behavior (Roy et al., 2021), and the static GLM-HMM recently applied to sensory decison-making behavior in fully trained animals (Ashwood et al., 2022; Bolkan et al., 2022). The dynamic GLM-HMM simultaneously characterizes discrete and continuous changes in choice policies, and thus is able to capture a wide range of non-stationary phenomena. For instance, the model can characterize individual differences in learning and task performance, disentangling whether an animal performs better due to increasing the weight on sensory cues or spending more time in an engaged state. The dynamic GLM-HMM is also well suited for simultaneously analyzing behavioral data during training and post-training epochs. This opens the door to novel analyses of behavior, since training data — which might be informative about post-training behavior — is often discarded.

The GLM-HMM was first used to model sensory decision-making during courtship in fruit flies (Calhoun et al., 2019). The authors identified three distinct sensorimotor strategies in male flies, each forming a distinct mapping between feedback cues from the female to the song mode produced by the male. Remarkably, they discovered a pair of neurons that drives switching between these multiple states or strategies. These findings show how GLM-HMM and its variants can be useful for characterizing behaviors across a wide range of subjects and experiments, even without any explicit task structure. Our dynamic extension could further reveal whether there are any experience-dependent changes occurring within these courtship strategies or in their usage, perhaps uncovering the effects of satiety or fatigue.

Recent work from Bruijns et al. (2023) introduced an extension of GLM-HMM known as the dynamic input-output infinite hidden semi-Markov model (diHMM), which provides a powerful model for dynamic state-dependent decision-making. The dynamic GLM-HMM and diHMM share similarities, with both models allowing for multiple decision-making strategies whose parameters vary smoothly over time. The generative model of the dynamic GLM-HMM could be considered a special case of diHMM, in which the number of total states is fixed and the durations of states follow a geometric distribution. Thus, diHMM is more flexible in its ability to capture more general patterns of state switching. However, the infinite model is harder to fit and interpret, as it requires sampling for inference and extensive post-hoc analysis to cluster states into “similar” types (Bruijns et al., 2023). By contrast, dynamic GLM-HMM has a more straightforward inference procedure, making it easier to fit to data and interpret.

Bruijns et al. (2023) applied clustering methods to the large number of states identified by their model, which led them to distinguish between three major training stages: “initial, biased behavior”, “partial, one-sided understanding of the task”, and “full understanding of the task”. This recapitulates previous findings showing that stimulus weights typically evolve away from 0 earlier on one side compared to the other (Roy et al., 2021). Importantly, Bruijns et al. (2023) also found that the majority of sessions were explained by a single state or strategy; this contradicts the findings of our model, which described most sessions as involving switches between multiple states. To examine this discrepancy, we performed a direct comparison between the dynamic GLM-HMM and diHMM fits on the same data. We found that held-out test data had significantly higher likelihood under the 3-state dynamic GLM-HMM than under the infinite semi-Markov model on most sessions (Supplemental Figure 3A). This suggests that our three-state model provides a better description of mouse behavior during learning than a model with a potentially unbounded number of states.

### Limitations and future directions

One extension of our work for obtaining a more fine-grained representation of learning would be to incorporate trial-to-trial changes in the weights. Although this would make our method more expressive, it does not constitute a serious limitation since previous work showed that largest changes in stimulus weights occur between sessions, after a “rest” period (Roy et al., 2021). Another future direction would be to infer during model fitting the hyperparameter *σ* dictating the rate of change of the weights across sessions. Our current procedure for finding *σ* is with cross-validation, which limits us to using the same *σ* across all states and features. An optimization over *σ* could tackle this limitation and use priors on the weights with different rates of change that depend on the specific state or input variable.

Although some tasks with non-stationary environments might require perpetual changes in the decision-making strategy to maximize reward (Piet et al., 2018), some of the changes in strategies that we’ve observed here lead to lower performance and less reward. It remains unknown why mice develop and use these disengaged or biased states. One possibility is that disengaged states reflect exploration (Dayan & Daw, 2008; Pisupati et al., 2021), whereas another possibility is that biased strategies require less motor, perceptual, or attentional effort (Hagura et al., 2017; Drugowitsch et al., 2012). New normative models are required to solve this problem.

To better understand why animals switch from one state to another, one future direction would be to model the state transitions using external variables, such as past sensory cues and accumulated reward. This would result in a more expressive structure for the hidden state transitions that would extend beyond the current Markovian rule (see Methods). To this end, recent work extended the standard GLM-HMM by modeling state transitions with a multinomial GLM using stimuli and reward histories (Mohammadi et al., 2024). Consistent with our findings that biased states align with the corresponding biased blocks in fully trained animals, they showed that past sensory stimuli govern the transition between the right-biased and left-biased states (Mohammadi et al., 2024).

Although our application of dynamic GLM-HMM here focused on learning, biases, and task engagement, our method could also identify other non-stationary phenomenona during decision-making, such as warm-up, satiety or fatigue. For instance, fatigue could be observed from a decrease in the likelihood of being in the engaged state towards the end of sessions. Since the current implementation of dynamic GLM-HMM can only be applied to two-alternative choice data, a future direction of our work consists of extending the output space to more than two categorical values or to continuous variables.

Lastly, the most important future direction would be to find neural correlates of these internal states and populations of neurons or regimes of activity that govern the switching between these states. Recent work made progress in this direction by finding three distinct populations in the median raphe nucleus that drive exploration, exploitation, and disengagement, respectively, during a multi-novel object interaction task (Ahmadlou et al., 2023). In all, there are many avenues for future research, yet we believe that we’ve developed a flexible and interpretable approach that is widely applicable to behavioral data.

## STAR ⋆ Methods

### Key resources table

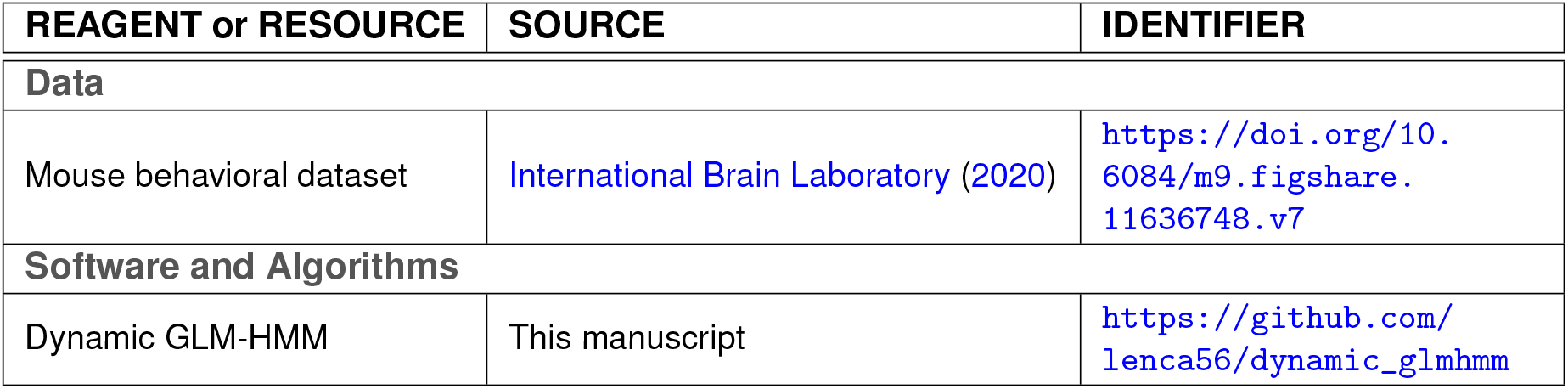

### Resource availability

#### Lead contact

Further information and requests for resources should be directed to the lead contact, Lenca I. Cuturela.

#### Materials availability

This study did not generate any new materials.

#### Data and code availability

- This paper analyzed publicly available behavioral data collected by the International Brain Laboratory groups (International Brain Laboratory, 2020), available at https://doi.org/10.6084/m9.figshare.11636748.v7. We used training and post-training data from 32 mice across 3 different labs (Witten, Churchland, and Angelaki labs).
- Our original code for fitting the dynamic GLM-HMM is distributed as a public GitHub repository at https://github.com/lenca56/dynamic_glmhmm.
- Any additional information required to reanalyze the data reported in this paper is available from the lead contact upon request.

### Experimental model and subject details

32 experimental subjects across 3 different labs were all female and male C57BL/6J mice aged 3-7 months, obtained from either Jackson Laboratory or Charles River. All procedures and experiments were carried out in accordance with the local laws and approval by the relevant institutions. This data was first reported in International Brain Laboratory et al. (2021).

### Method Details

#### Dynamic GLM-HMM

The dynamic GLM-HMM characterizes time-varying behavioral strategies by allowing the transition matrix *P*^*s*^ and the state-dependent GLM weights 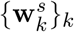 to vary per-session *s* for each state *k*. An animal’s binary choice 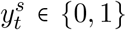 at trial *t* in session *s* (where 0 and 1 correspond to leftward and rightward choices, respectively) is dependent on the current state via a latent variable 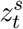. More precisely, the probability of a choice 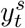 is a linear combination of the task covariate vector 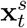 and the state-specific GLM weights passed through a logit function:

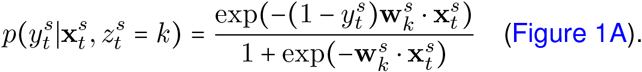

The *D*-element vector 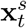 contains task variables that are thought to affect an animal’s choice, such as the signed stimulus contrast, a bias term, the previous rewarded choice, and the previous choice. The larger the absolute value of a specific weight, the more the animal relies on that corresponding variable when making a choice. The sign of the weight represents the direction of this effect towards one side compared to the other.

The *K × K* session-specific transition matrix *P*^*s*^ defines the probability of transitioning between discrete states in consecutive trials during session *s*:

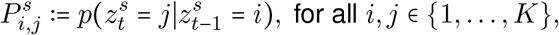

where *K* is the total number of hidden states. Note that the transition between states has the Markov property, meaning that the current state only depends on the previous one. Moreover, each row of the transition matrix sums up to 1 and all its entries are non-negative. The first latent state for each session *s* is drawn uniformly by 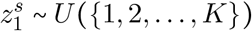.

The weights vary across sessions according to a Gaussian prior,

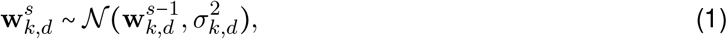

where *σ*_*k,d*_ is a positive hyperparameter that determines the variability of the weights across sessions for state *k* and task variable *d*. A larger *σ*_*k,d*_ implies a larger session-to-session change for the weights 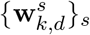.

Each row of the transition matrix varies across sessions according to a Dirichlet prior,

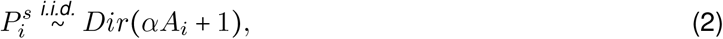

where *A* is the inferred global transition matrix, and *α* is a non-negative scalar hyperparameter that governs the intensity of the Dirichlet distribution. When *α* is large, the session-specific transition matrices *P* ^*s*^ are all close to the global transition matrix *A*. Note that this prior was chosen for closed-form optimization and does not impose temporal continuity or smoothness in the transition matrix across session.

In the limit case when all *σ*_*k,d*_ ↓0 and *α* → ∞, the transition matrices *P*^*s*^ are all equal to the global matrix *A*, and the weights are all constant across sessions, hence our method becomes equivalent to the standard GLM-HMM (Bolkan et al., 2022).

#### Inference of dynamic GLM-HMM parameters

##### MAP estimation

The goal of the dynamic GLM-HMM inference procedure is to fit the model parameters using Maximum A Posteriori (MAP) estimation, given a set of fixed hyperparameters *K*, {*σ*_*k,d*_}, *α*. This entails learning the session-specific transition matrices *P*^*s*^ and weights 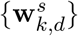 collectively labeled as *θ*. This was implemented with a variation of the Expectation-Maximization (EM) algorithm, similarly to other previous adaptations for fitting input-output HMMs (Bengio & Frasconi, 1994; Escola et al., 2011; Calhoun et al., 2019; Ashwood et al., 2022).

The EM algorithm maximizes the log-likelihood of the data and can be extended to maximize the log-posterior of the parameters given the data, where the data is denoted by 𝒟 = {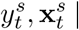 for all trials *t* and sessions *s*}. Using Bayes’ rule, the log-posterior can be computed up to an irrelevant constant by:

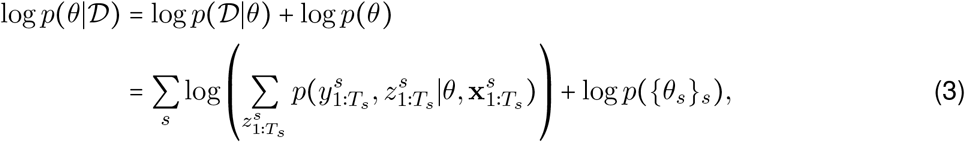

where the sum over 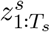 is over all possible latent states in session *s*.

The log-prior on the parameters for a session *s* (given that the parameters from all other sessions are fixed) can be computed using Equation 1 and 2:

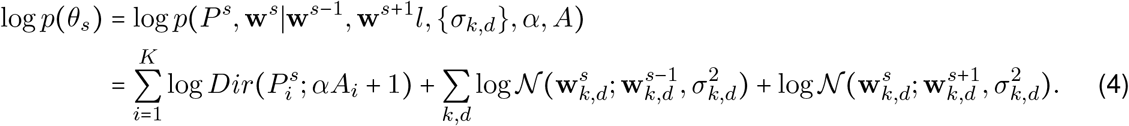

##### Expectation-Maximization algorithm

The EM algorithm maximizes the log-posterior in Equation 3 by alternating between an Expectation step (E-step) and a Maximization step (M-step) for multiple iterations (Bishop, 2006; Bengio & Frasconi, 1994). The sum in Equation 3 is not directly optimized since it involves an exponential number of terms 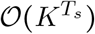 that arise due to the summation over all possible latents 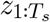 in a session *s*. Instead, the EM algorithm provides an efficient way around that by maximizing at every M-step w.r.t to *θ* a quantity *Q (θ, θ*^*old*^ ), which is known as the “Expected Complete-Data Log-Likelihood” (ECLL). The ECLL, together with the prior on the parameters, provides a lower bound on the log-posterior:

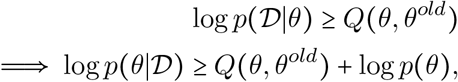

where *θ*^*old*^ are the parameters from the previous E-M iteration and *θ* are the parameters of the current iteration that are being optimized (Bishop, 2006; Bengio & Frasconi, 1994). Thus, maximizing at every M-step the ECLL together with the prior guarantees that the log-posterior increases as a function of the parameters *θ*.

In particular, our inference procedure performs an E-step and an M-step for each session (in which the parameters from all other sessions are fixed) and iterates across all sessions multiple times until all parameters converge to a local or global optimum.

##### Expectation step (Forward-Backward algorithm)

The E-step for a session *s* computes the session-specific ECLL represented through the auxiliary function,

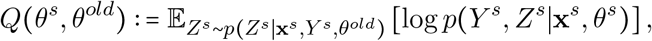

where *θ*^*old*^ are all the parameters from the previous E-M iteration, *θ*^*s*^ are the parameters of the current iteration, 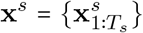 are the inputs or task variables, 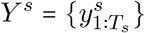 are the outputs (animals’ choices), and 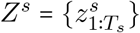 are the latents for session *s*.

###### Notation

In order to efficiently compute *Q* (*θ*^*s*^, *θ*^*old*^) at the E-step, we will introduce the following notations. The observation probabilities of the choices are denoted by

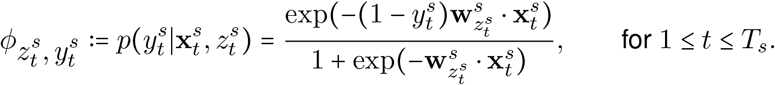

The forward conditional probability of the latents is

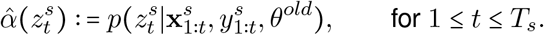

The backward conditional probability of the latents is

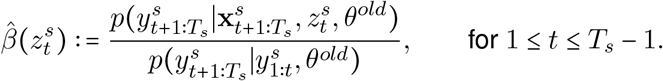

The scaling factor is

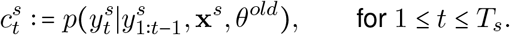

The marginal posterior probability of the latents is

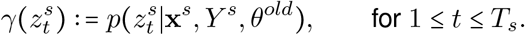

Lastly, the joint posterior probability of two successive latents is

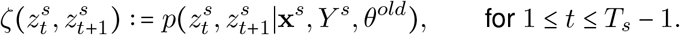

For every session *s*, all probabilities are conditioned on 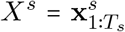, which we’ll omit writing from now on.

The expected complete data log-likelihood for a session *s* can be rewritten as follows:

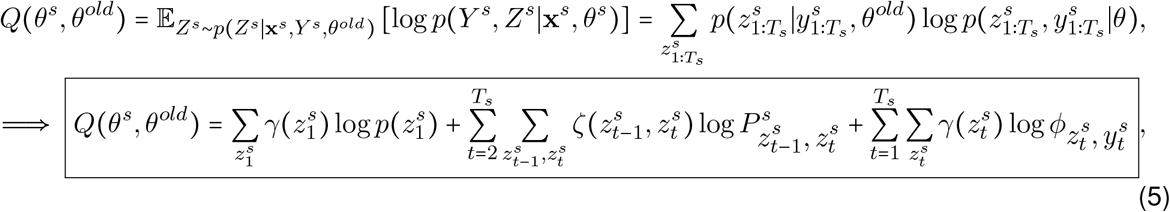

where *ϕ* depends on the current weights 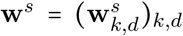 that are to be optimized together with the transition matrix *P*^*s*^, whereas 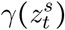 and 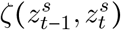 are computed using the previous parameters *θ*^*old*^. We compute the posterior probabilities of the latents 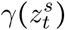 and 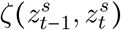 as follows, where *θ*^*old*^ is implicitly conditioned on everywhere:

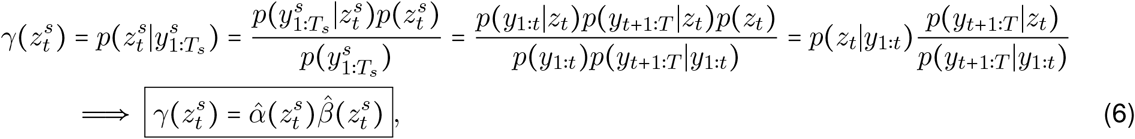

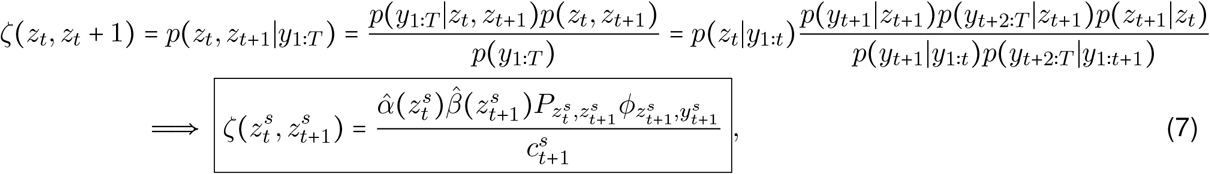

which depend on the forward and backward probabilities of the latents 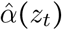 and 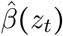, respectively.

###### Forward-Backward algorithm

We use the forward-backward algorithm, a two-step message passing algorithm (Bishop, 2006), in which forward and backward probabilities propagate in time, to compute recursively 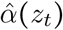 and 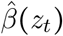 as follows, where *θ*^*old*^ is implicitly conditioned on everywhere:

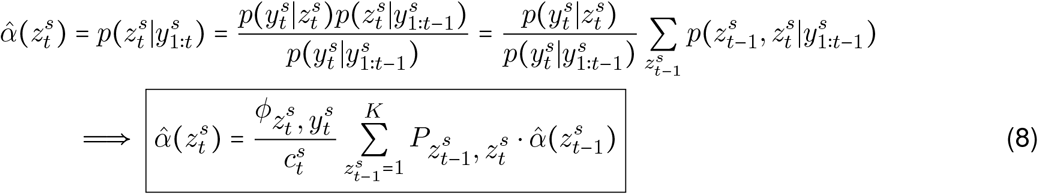

where 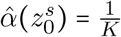 represents the prior on the latent states before having seen any data.

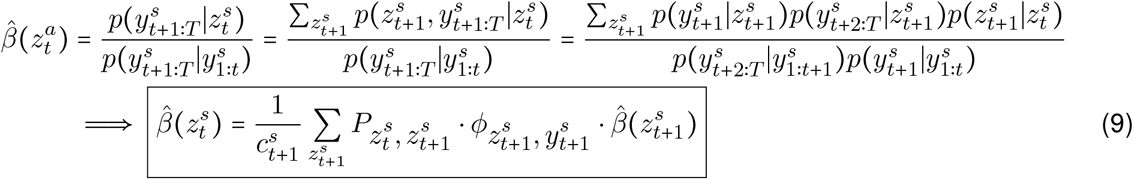

where 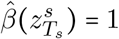.

##### Maximization step

After running the forward-backward algorithm, Equation 5, 6, 7, 8, 9 together give the ECLL for each session. Then, we maximize with respect to the session-specific parameters, *θ*^*s*^ ={ **w**^*s*^, *P*^*s*^ }, the persession ECLL together with the log-prior, which is given by Equation 4. Note that the ECLL written as in Equation 5 contains weight terms **w**^*s*^ (in the observation probabilities *ϕ*) and transition matrix terms *P* ^*s*^ that are added together. This means that optimization is done independently for *P* ^*s*^ and **w**^*s*^.

Firstly, we maximize with respect to the per-session transition matrix *P* ^*s*^ the function

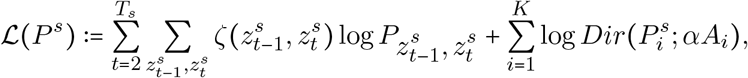

which results in the unique closed-form update of the transition matrix:

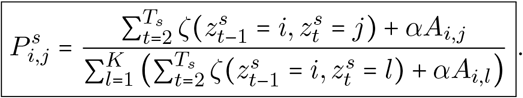

To do so, we use the Lagrange multipliers method since each row of the transition matrix is constrained to sum up to 1. Note that the term containing 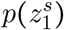 in Equation 5 is not optimized since the prior on the first latent state of any session is the uniform distribution on all the possible states,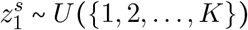.

Secondly, we maximize with respect to the per-session weights **w**^*s*^ the function

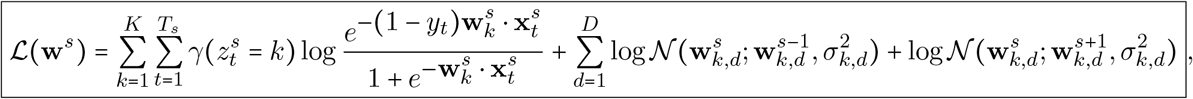

where for the first session, *s* = 1, we only have one prior term coming from the next session, and for the last session, *s* = *S*, we only have one prior term coming from the previous session. Since there is no closed-form update for the weights, we use scipy.optimize.minimize with a quasi-Newton optimization algorithm called ‘BFGS’ to find the weights **w**^*s*^ that (locally) maximize the above function (Virtanen et al., 2020).

##### Cross-validation and model performance

To decide the number of states in the model *K*, and the hyperparameters *σ* and *α* dictating the variability of the weights and transition matrix, respectively, we implemented a 5-fold cross-validation procedure to find the best-fitting models. We set *σ*_*k,d*_ = *σ* for all states and features to avoid very long fitting times.

Model performance is quantified through the marginal log-likelihood of test data,L_*test*_(*θ*_*train*_):= log *p*(*Y*_*test*_|**x**_*test*_, *θ*_*train*_),given the parameters obtained by fitting the model to the training set, *θ*_*train*_ = { *P*_*train*_, **w**_*train*_ }. For each session, 80% of trials (randomly selected in consecutive blocks of 10) belong in the train set, whereas the other 20 % of trials belong to the test set. The test log-likelihood for a model is then obtained by averaging the test log-likelihoods across the 5 different splits. We normalize the test log-likelihood for each session by the total number of trials during that session, and then multiply it with the average session length to get the “average test log-like (session)” used in Figure 2B, 3E, 4A,B, 5A.

We can compute the marginal log-likelihood for any set of parameters *θ* during the forward pass of the forward-backward algorithm:

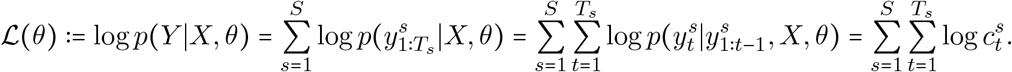

The hyperparameters that led to highest model performance across animals are σ ≈3.2 and σ ≈2. Note that we chose *K* = 3 for our final models since there was not a substantial difference in performance when extending to *K* = 4 or *K* = 5 states.

##### Multi-stage fitting procedure and initialization

When fitting our method to data from each animal individually, we initialize the parameters from the global standard GLM-HMM fitted parameters across all mice. This approach allows us to avoid cumbersome fitting procedures that require many random initialization for the time-varying parameters. Moreover, this choice of initialization is consistent with the idea that individual animal fits should be relatively close to the global average fit.

The obtain the static, global transition matrix *A*, we first fit the dynamic GLM-HMM with a constant transition matrix across sessions and time-varying weights, with the initialization described above. We then use the fit transition matrix *A* for the dynamic prior on the time-varying transition matrix as in Equation 2.

Our multi-stage fitting and initialization procedure is summarized below:

**For** each number of states *K* ∈ {1, 2, 3, 4, 5 }do:

Fit standard GLM-HMM to data from all animals together:

**For** init ∈ {1, 2, … , 20}

Initialize K-state standard GLM-HMM from noisy parameters

Run EM algorithm until parameter convergence

Find K-state model with parameters *W*_′_ and *P*_′_ that performs best on the test set

Fit dynamic GLM-HMM with dynamic weights and static transition matrix for each animal:

**For** *σ* ∈ { 10 ^™3^, 10 ^™2.5^, 10 ^™2^, ⋯, 10^2.5^, 10^3^}:

Initialize dynamic GLM-HMM using fitted static global parameters *W′* and *P*^′^

Run EM algorithm until parameter convergence

Find *σ′* that corresponds to best performance on the test set

Let recovered parameters from best-fitting model be *W*^′′^ and *A*

Fit dynamic GLM-HMM with dynamic weights and dynamic transition matrix for each animal:

**For** *α* ∈ { 2 *10 ^™1^, 2 *10 ^™0.5^, ⋯ 2 * 10^5^}:

Initialize K-state dynamic GLM-HMM from previous fit parameters *W* ^′′^ and *A*

Use previously found hyperparameter *σ′* and *A* for the dynamic priors

Run EM algorithm until parameter convergence

Find *α′* that corresponds to best performance on the test set

Find number of states *K* that corresponds to best performance on the test set

##### Significance testing

For each animal individually, we used a one-sided paired t-test for comparing task-accuracy in state 1 to average task-accuracy in states 2 and 3 across sessions. We obtained significantly better performance in state 1 for each animal, with the following p-values for all animal rounded up to 3 decimals: (0., 0., 0., 0., 0., 0., 0., 0., 0., 0., 0., 0. 0., 0., 0., 0., 0., 0., 0., 0.002, 0., 0., 0., 0., 0., 0., 0., 0., 0., 0., 0., 0.014).

Across all animals, we used a one-sided paired t-test to compare time spent in state 1 between consecutive blocks of 5 sessions. We found that animals spend significantly less time in state 1 during sessions 0-5 vs 5-10 (p=0.009) and during sessions 5-10 vs 10-15 (p=0.013). The animals did not spend significantly different time in state 1 between sessions 10-15 and 15-20 (p=0.407).

We performed a one-sided paired t-test to compare the log-likelihood under our 3-state dynamic GLM-HMM and one under the dynamic infinite semi-Markov model from Bruijns et al. (2023). The two models fits used the same train-test splits of the data across 20 different IBL animals. We found that 3-state dynamic GLM-HMM performs significantly better on the majority of sessions during learning Supplemental Figure 3A, with the following p-values across sessions rounded up to 3 decimals: (0.978, 0.0, 0.0, 0.0, 0.0, 0.0, 0.0, 0.0, 0.0, 0.005, 0.0, 0.0, 0.0, 0.0, 0.001, 0.003, 0.013, 0.001, 0.167, 0.024, 0.42, 0.086, 0.016, 0.014, 0.016, 0.037, 0.056, 0.005, 0.141, 0.018).

## Acknowledgements

We are grateful to Peter Dayan, Liam Paninski, Sebastian Bruijns, and Victor Geadah for critical feedback on the manuscript, and to Sebastian Bruijns for help in performing the model comparisons shown in Supplemental Figure 3. This work was supported by grants from the Simons Collaboration on the Global Brain (SCGB AWD543027), the NIH BRAIN initiative (9R01DA056404-04), and a U19 NIHNINDS BRAIN Initiative Award ((U19NS123716).

## Supplemental information

### Dynamic GLM-HMM recovers true parameters on simulated data

We validated our method by testing its ability to recover the true parameters from simulated data (Supplemental Figure 1). Firstly, we fit the dynamic GLM-HMM to simulated data in which the true transition matrix is constant across sessions. This will provide an estimate for the global transition matrix *A* that we can then use for fitting a time-varying transition matrix *P*^*s*^ with a dynamic prior that depends on *A*.

We fit multiple models with different number of states and values of *σ*, which dictates the rate of change across all the weights (*σ*_*k,d =*_ *σ*). We chose to have the same rate of change across all states and features to avoid a slow fitting procedure and we acknowledge that *σ* will represent a compromise between the variability of the drifting weights in state 1 and the static weights in state 2. Using cross-validation, we find that the highest log-likelihoods on the test set are attained for *σ* ≈ 0.1 across all models with 1, 2, and 3 states, respectively (Supplemental Figure 1A). As expected, the standard GLM-HMM is outperformed by a dynamic GLM-HMM with *σ* > 0 for all models since the true weights are not static. Moreover, the one-state model performs the worst across all values of *σ* and is unable to capture the full variability of the data, whereas the two-state and three-state models attain similar test log-likelihood at optimal *σ* ≈ 0.1. Since two discrete states are sufficient to reach highest model performance, our method can recover the true number of latent states. Interestingly, the standard GLM-HMM performs substantially better for the three-state model compared to the two-state one, which results in a mismatch between the inferred number of states and the true one. Since the standard GLM-HMM attempts to capture dynamic weights through a set of constant ones, we posit that the method will often overestimate the number of states when animals use behavioral strategies that evolve over time. This is a significant limitation of the standard GLM-HMM that our method improves upon.

**Supplemental Figure 1:**
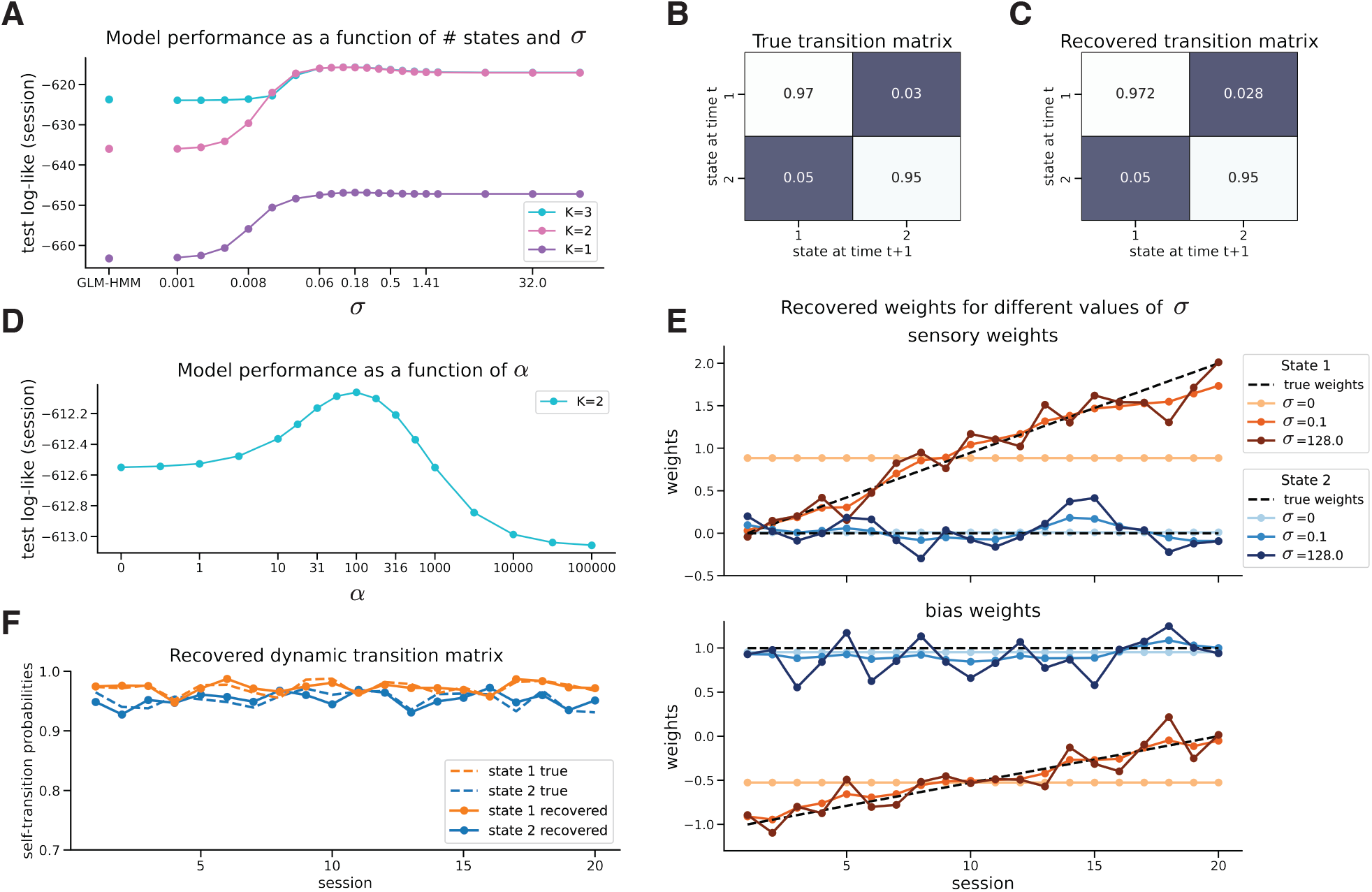
Dynamic GLM-HMM recovers true parameters on simulated data. **A**. Test log-likelihood (average per session) as a function of hyperparameter *σ* for dynamic GLM-HMM models with 1, 2, and 3 states, respectively. **B**. True constant transition matrix. **C**. Recovered static transition matrix for best-fitting two-state model with *σ* = 0.1. **D**. Test log-likelihood (session) as a function of transition matrix hyperparameter *α* for two-state dynamic GLM-HMM with time-varying parameters. **E**. True and fit sensory weights (top) and bias weights (bottom) for the two-state models with different values of *σ*. **F**. True and recovered dynamic transition matrix for the best-fitting 2-state model with *α* = 100.

The recovered static transition matrix for the two-state model closely matches the true one (Supplemental Figure 1B,C). The recovered weights for the best-fitting two-state model (*σ* ≈ 0.1) also agree with the true ones, hence validating our inference procedure (Supplemental Figure 1E). Note that *σ* smaller than the optimal one leads to an underestimation of the variance in the recovered weights, whereas a *σ* larger than the optimal one overestimates the variance of the weight and results in over-fitting (Supplemental Figure 1E). At *σ* = 0, dynamic GLM-HMM uses constant weights across sessions, being equivalent to a standard GLM-HMM. Note that very large values of *σ* result in recovered weights close to the maximum likelihood estimator for each session since the prior has little to no effect (Bishop, 2006). In either extreme, the recovered weights do not match the true weights, emphasizing the importance of choosing an appropriate drifting parameter *σ*.

**Supplemental Figure 2:**
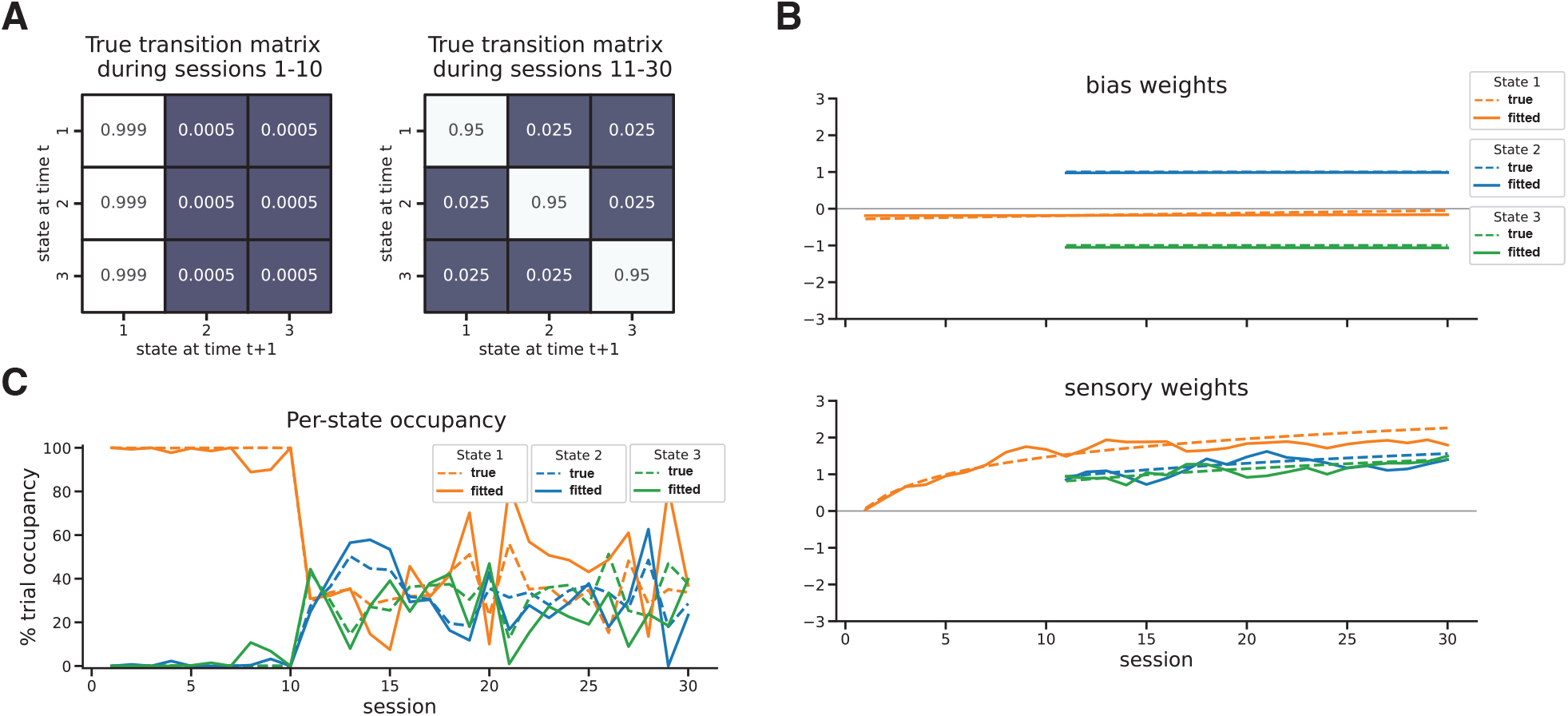
Dynamic GLM-HMM can flexibly use different number of states across sessions. **A**. True 3 x 3 transition matrices used for simulating data during first 10 sessions (left) and after the first 10 sessions (right). **B**. True vs fitted bias weights (top) and sensory weights (bottom) for the 3 states as a function of session number. **C**. % trials spent in each state as a function of session number for the true and fitted parameters.

Secondly, we apply our method to simulated data when the true transition matrix is non-stationary. For the dynamic prior on the session-specific transition matrices, we use the above estimate of the global transition matrix *A* (Supplemental Figure 1C). By cross-validation, we find the optimal intensity hyperparameter *α* to be approximately 100, which aligns with the true *α* used for simulating the true dynamic transition matrix (Supplemental Figure 1D). Importantly, the recovered dynamic transition matrix closely matches the true one, confirming that our method can recover all dynamic parameters well (Supplemental Figure 1F).

Last but not least, we apply our method to a different simulated dataset in which an imagined animal only employs a single unbiased strategy during the first 10 sessions (Supplemental Figure 2). The transition matrix initially has probabilities close to 1 of transitioning to state 1 from every state, which corresponds to 100% of trials spent in state 1 during the first 10 sessions (Supplemental Figure 2A,C). Later on, the transition matrix changes drastically to have high probabilities on all the diagonal elements, which correspond to long blocks of consecutive trials and a similar number of trials spent in each state (Supplemental Figure 2A,C). Given the same amount of trials as in a real IBL experiment, we are able to recover the true parameters and state occupancies across sessions (Supplemental Figure 2B,C).

### Comparison between dynamic GLM-HMM and diHMM

We performed a comparison of the log-likelihood under our 3-state dynamic GLM-HMM and the dynamic infinite semi-Markov model from Bruijns et al. (2023). The two models fits used the same train-test splits of the data across 20 different IBL animals. Note that Bruijns et al. (2023) use distinct inputs for the left and right sensory cues in order to characterize differences in performance on one side compared to the other, whereas we use the signed stimulus contrast that captures the performance on both sides together. We find that 3-state dynamic GLM-HMM performs significantly better (*p* < 0.05) on the majority of sessions during learning (Supplemental Figure 3A).

**Supplemental Figure 3:**
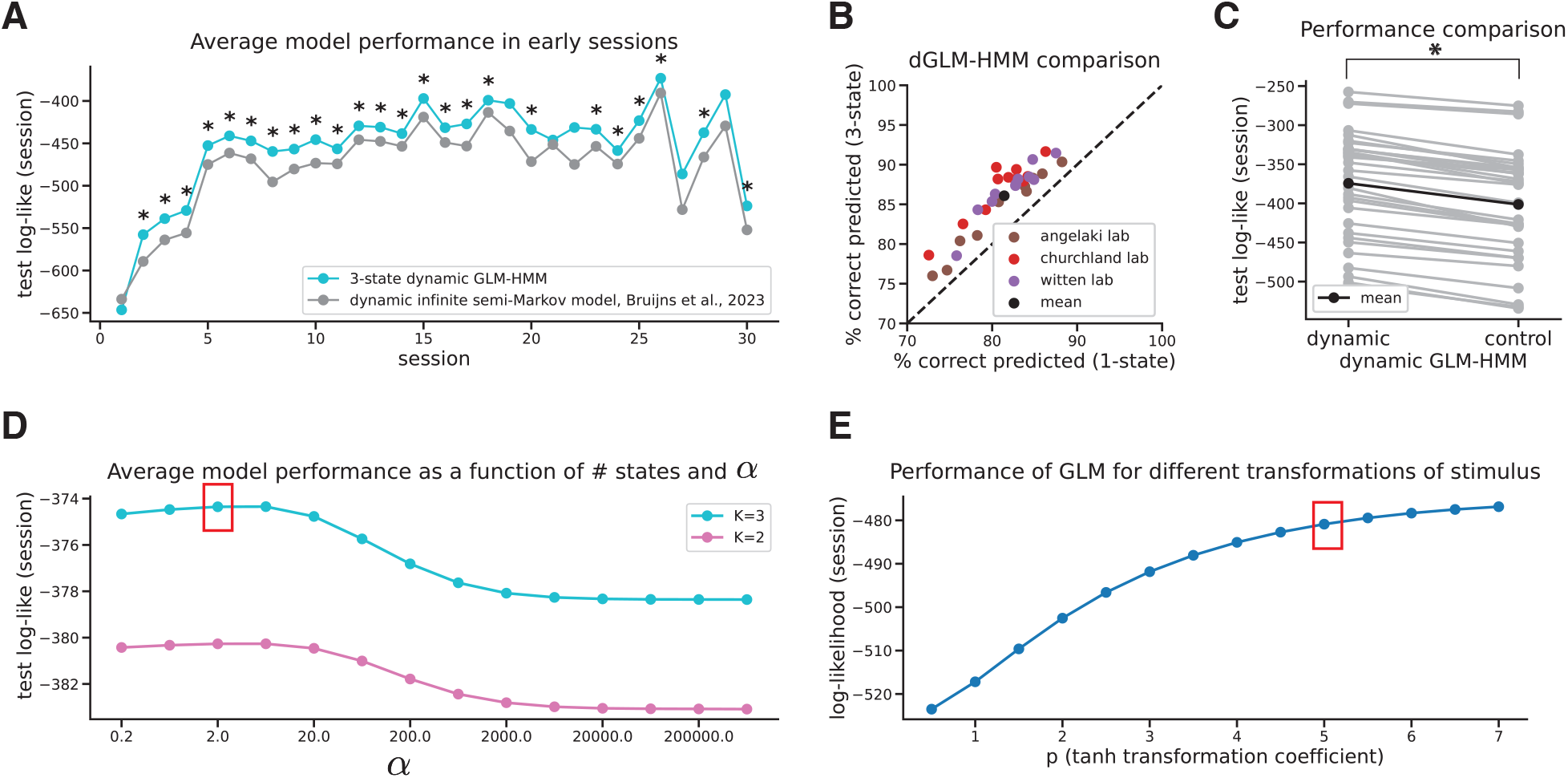
Dynamic 3-state GLM-HMM achieves better performance. **A**. Average test log-likelihood across 20 animals as a function of session number for our best-fitting 3-state dynamic model compared to best fits from Bruijns et al. (2023). A star for any session means that our model performed significantly better on that session (paired t-test, *p* < 0.05). **B**. % correct prediction of animals’ choices for best 1-state dynamic GLM-HMM vs 3-state one, where each point represents a different animal (the mean is shown in black) **C**. Log-likelihood (per session), averaged across sessions and animals (in black) or individually for each animal (in gray) for our best-fitting 3-state dynamic model and a control model, in which all parameters are identical to the best-fitting ones except the stimulus weights in biased states 2 and 3, which were enforced to be static and are set to the mean of the corresponding time-varying weights in the fully dynamic model. The 3-state dynamic model performed significantly better than the control (paired t-test, *p* < 0.05). **D**. Δ test log-likelihood (averaged per session) as a function of hyperparameter *α* for models with 2 and 3 states, respectively, averaged across all animals, and compared to 1 state model performance. The red rectangle encloses the best-fitting model (*K* = 3 and *α* = 2). **E**. Average log-likelihood per trial for classic GLM, fit to data across all animals, as a function of tanh transformation coefficient p. The red rectangle represents the choice of *p* = 5 for all other analysis in the paper.

### Trial history is not an important regressor for choice behavior

We modeled animals’ choices using four task inputs: the signed stimulus contrast (positive for the right side and negative for the left side), an offset or bias, the animal’s choice on the previous trial, and the rewarded side of the previous trial. A large weight on the animal’s previous choice reflects a strategy termed “perseveration”, in which the animal tends to repeat the previous choice irrespective of the rewarded side. A large weight on the rewarded side of the previous trial reflects a strategy known as “win-stay, lose-switch”, in which the animal repeats a choice if it was previously rewarded and switches otherwise.

For the example mouse, the history-dependent weights are substantially smaller than the bias and stimulus weights (Supplemental Figure 4A). On average across animals, both the previous choice and previous correct side weights are consistently close to 0 across all states, suggesting that these inputs are not important in determining the state-dependent strategies (Supplemental Figure 4B).

**Supplemental Figure 4:**
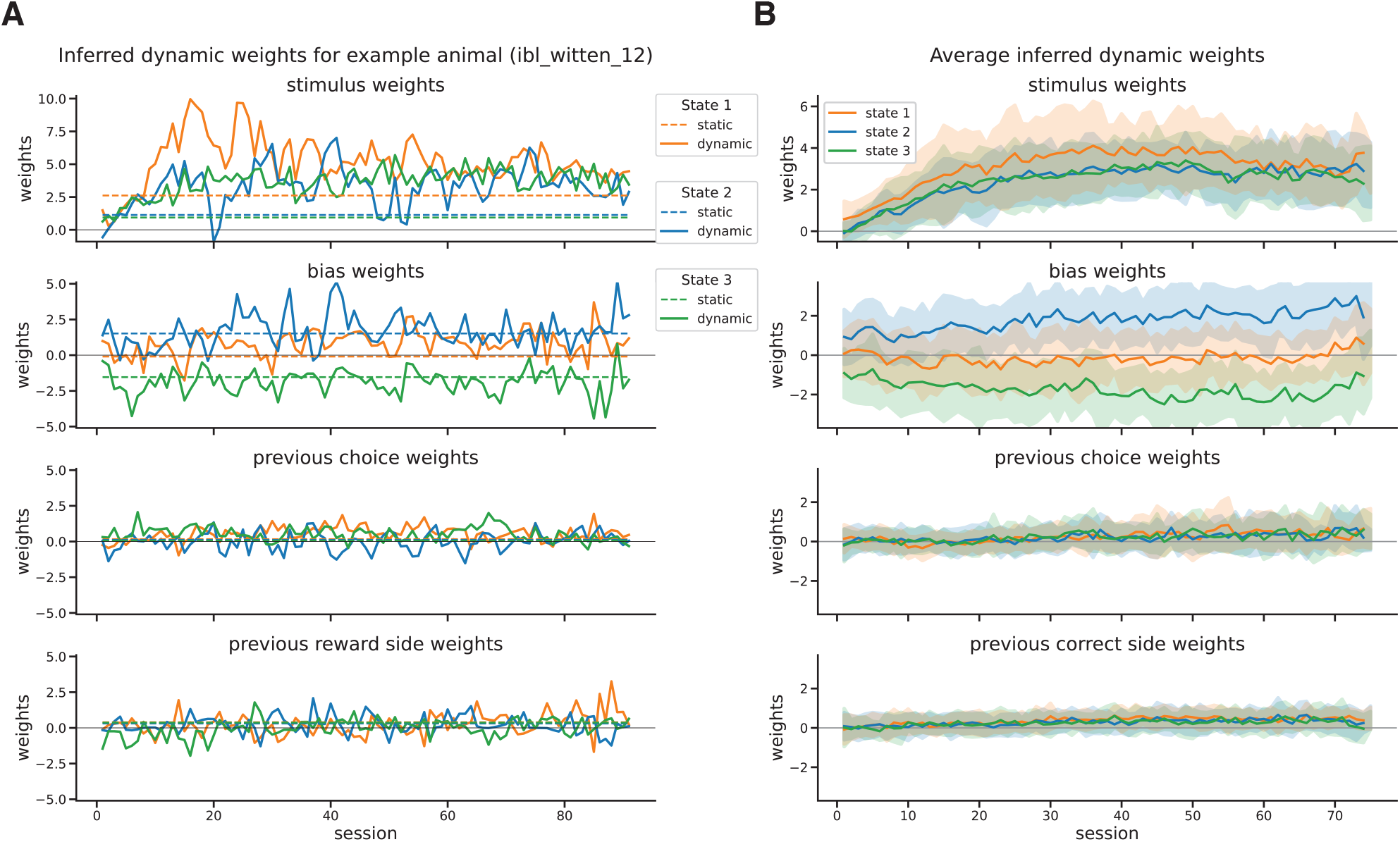
Trial history is not an important regressor for choice behavior. **A**. Inferred dynamic GLM weights for the best fitting 3-state model (*σ* = 1) for four task variables: signed stimulus contrast, bias, previous choice, and previous rewarded side. Inferred weights for the standard GLM-HMM are shown with dashed lines. **B**. Average inferred dynamic GLM weights for the best 3-state model (*σ* = 1) for the four task variables. Individual weights are shown in the background for each animal.

